# Astrocyte-Derived Exosomes Regulate Sperm miR-34c Levels to Mediate the Transgenerational Effects of Paternal Chronic Social Instability Stress

**DOI:** 10.1101/2023.04.21.537854

**Authors:** Alexandre Champroux, Mitra Sadat-Shirazi, Xuan Chen, Jonathan Hacker, Yongjie Yang, Larry A. Feig

## Abstract

The effects of chronically stressing male mice can be transmitted across generations by stress-specific changes in their sperm miRNA content that induce stress-specific phenotypes in their offspring. But how each stress paradigm alters the levels of distinct sets of sperm miRNAs is not known. We showed previously that exposure of male mice to chronic social instability (CSI) stress results in elevated anxiety and reduced sociability specifically in their female offspring across multiple generations because it reduces miR-34c levels in sperm of stressed males and their unstressed male offspring. Here we describe evidence that **a**strocyte-derived **exos**omes (**A-Exos**) carrying miR-34c mediate how CSI stress has this transgenerational effect on sperm. We found that CSI stress decreases miR-34c carried by A-Exos in the prefrontal cortex and amygdala, as well as in the blood of males. Importantly, miR-34c A-Exos levels are also reduced in these tissues in their F1 male offspring, who despite not being exposed to stress exhibit reduced sperm miR-34c levels and transmit the same stress-associated traits to their male and female offspring. Furthermore, restoring A-Exos miR-34c content in the blood of CSI-stressed males by intravenous injection of miR-34c-containing A-Exos restores miR-34c levels in their sperm. These findings reveal an unexpected role for A-Exos in maintaining sperm miR-34c levels by a process that when suppressed by CSI stress mediates this example of transgenerational epigenetic inheritance.

## 1 INTRODUCTION

The concept of transgenerational epigenetic inheritance, where traits acquired from environmental exposures are transmitted to subsequent generations through epigenetic modifications in germ cells, is well established in organisms such as *C. elegans*, plants, and fruit flies[1, 2]. Growing evidence supports this concept in rodents, demonstrating that environmental factors like chemicals [3], drugs [4], diet [5], enriched environment[6], and chronic stress [7] lead to altered phenotypes in offspring across multiple generations. For male exposures, epigenetic alterations in sperm, including altered levels of miRNAs, have been implicated. Human epidemiological studies also align with this concept [8], including summary statistics for eleven neuropsychiatric disorders that imply that the fraction of the heritability of these maladies not accounted for by classical genetics may be due to epigenetic inheritance [9].

A striking yet unexplained aspect of this process in mice is how different chronic stress paradigms alter the content of distinct sets of sperm miRNAs, leading to different behavioral effects in offspring. For instance, Gapp et al [10] found that male mice exposed to unpredictable maternal separation at birth combined with unpredictable maternal stress (**MUS**) exhibited increased levels of several sperm miRNAs, including miR-375, miR-200b, miR-672, and miR-466. When all sperm RNAs from these stressed mice were injected into control zygotes, the resulting male offspring displayed heightened anxiety and depression, mirroring the behavioral changes seen in the offspring of stressed males [10]. In contrast, Rogers et al [11] showed that chronic variable stress (**CVS**) in adult males elevated levels of 8 different sperm miRNAs (plus miR-375 in common). Injection of these 9 miRNAs into control zygotes also replicated the observed effects in the offspring of stressed males. No basal behavior changes were detected, but instead a blunted HPA axis response to stress was found in both males and females.

We demonstrated that exposing adolescent male mice to chronic social instability (**CSI**) stress, characterized by introducing a novel mouse twice a week for 7 weeks, results in elevated anxiety and impaired sociability specifically in female offspring for three generations through the paternal lineage [12]. CSI stress induces a decrease in sperm levels of miR-34/449 family members [13] and an increase in miR-409-3p [14] not only in directly stressed males but also in the sperm of their F1 male offspring. These F1 males, who were not exposed to stress, still transmit the same stress-associated traits to their male and female offspring.

Furthermore, we found that the reduced levels of miR-34/449 in the sperm of stressed males are passed on to **P**re**I**mplantation **E**mbryos (**PIEs**) upon fertilization [13]. We then showed that this decrease in miR-34/449 levels in PIEs is critical for transmission of stress-related traits to offspring, since rescue of suppressed levels of these miRNAs reversed the behavioral phenotypes in female offspring. It also suppressed the appearance of reduced levels of miR-34/449 in sperm of male offspring that are responsible for making this phenomenon transgenerational, not just intergenerational [15].

Elevated stress hormones might contribute to the alterations in sperm miRNA profiles induced by chronic stress, since injection of glucocorticoids can alter sperm miRNA levels. However, the miRNAs affected did not match any of those described above [16], implying that a more targeted signaling mechanism is involved. Moreover, changes in blood metabolites have been linked to stress-induced changes in spermatogonia and offspring metabolism, but the source and role of these metabolites in transmitting behavioral change to progeny remain unclear [17]. In this study, we provide evidence that, for CSI stress, a more specific signaling mechanism involves reduced miR-34c in astrocyte-derived exosomes. These exosomes, originating from a subset of brain regions known to respond to stress, are transported via the blood to regulate its level in sperm, and thus mediate this example of transgenerational epigenetic inheritance.

## 2 METHODS

### 2.1 Animals and housing

All mice in this study were of the CD-1 strain obtained from Charles River Laboratories. Males used for CSI stress began the protocol at 28 days postnatal age, and control females used for breeding were 8 weeks of age. All animals were housed in temperature, humidity, and light-controlled (14hr on/10hr off LD cycle) rooms in a fully staffed dedicated animal core facility led by on-call veterinarians at all hours. Food and water were provided *ad libitum*. All procedures and protocols involving these mice were approved by the Institutional Animal Care and Use Committee of the Tufts University School of Medicine, Boston, MA.

### 2.2 Chronic social instability (CSI) stress

CSI protocol was conducted as described previously^6^. Briefly, the composition of each mouse cage (four mice per cage, 28 days old) was randomly shuffled twice per week for 7 weeks, such that each mouse was housed with three new mice in a fresh, clean cage. Control mice were housed with the same four mice per cage for the duration of the protocol. Any mice involved in physical interactions associated with this stressful condition are removed. After 7 weeks, mice were housed in pairs with a cage mate from the final cage change and left for 2 weeks to remove the acute effects of the final change. Mice were either sacrificed for sperm, blood, and brain collection or mated with random control female mice overnight to generate “F1” animals. We have previously shown that stressed males can be mated multiple times and still transmit stress phenotypes to their offspring [12], so male mice were used for mating or sperm collection as needed.

### 2.3 Mouse sperm collection

Mature, motile mouse sperm (from the cauda epididymis) were isolated via the swim-up method [18]. Briefly, male F0 and F1 mice were anesthetized under isoflurane and sacrificed via cervical dislocation. The caudal epididymis and vas deferens were dissected bilaterally and placed in 1mL of warm (37°C) M16 medium (Sigma-Aldrich, M7292) in a small Petri dish. Under a dissection microscope, sperm were manually expressed from the vas deferens using fine forceps, and the epididymis was cut several times before incubating at 37°C for 15 min to allow mature sperm to swim out, then large pieces of tissue were removed. The remainder of the extraction took place in a 37°C warm room. The sperm-containing media was centrifuged at 3,000 RPM for 8min, the supernatant was withdrawn and discarded, and 400μL of fresh, warm M16 medium was carefully placed on top of the pellet. The tubes were then allowed to rest at a 45° angle for 45 min to allow the motile sperm to swim-up out of the pellet into the fresh medium. The supernatant containing the mature sperm was carefully withdrawn and centrifuged again for 5 min at 3,000 RPM to pellet the motile sperm. The supernatant was withdrawn and discarded, and the pellet was frozen on dry ice for later processing.

Caput sperm were collected as described[19].Two small incisions were made at the proximal end of caput and using a 26G needle holes were made in the rest of the tissue to let the caput epididymal fluid ooze out. Sperm-containing media was incubated for 15 minutes at 37°C, then transferred to a fresh tube and the sperm-free epididymis tissues were directly frozen in liquid nitrogen and stored at -80°C. The sperm were incubated for another 15 minutes at 37°C, then sperm were collected by centrifugation at 2,000 x g for 2 minutes, followed by a 1X PBS wash, and a second wash with somatic cell lysis buffer (low SLB buffer, 0.01% SDS, 0.005% Triton X-100 in PBS) for 10 minutes on ice to eliminate somatic cell contamination. Somatic cell lysis buffer treated sperm were collected by centrifugation at 3,000xg for 5 minutes, and finally washed with 1X PBS before freezing down.

### 2.4 Epididymosome isolation by ultracentrifugation and density-gradient

After pelleting cauda sperm following the swim-up protocol in M16 medium (Sigma, M7167), the supernatant was centrifuged at 2,000xg for 10min, 10,000xg for 30min and then ultracentrifuged at 120,000xg at 4°C for 2h. The epididymosomal pellet was then washed in PBS 1X at 4°C and ultracentrifuged at 120,000xg at 4°C for 2h. The resulting pellet was resuspended in 50µl of PBS [20].

### 2.5 Submandibular blood collection

The submandibular blood was collected with K2E (Bd microtainer) from the submandibular vein using an animal lancet (size 5.5 mm, Goldenrod, Medipoint). Then the blood was centrifuged at 2,000xg, for 15min at 4°C before frozen for RNA extraction.

### 2.6 Isolation of exosomes from plasma

Blood was collected by cardiac puncture of the heart of the mice before sacrifice. Then the blood was centrifuged at 2,000xg for 15min at 4°C before frozen. Total exosomes from the plasma were purified using the Total exosomes isolation kit (Invitrogen, #4484450). Briefly, plasma was thawed at 36°C and then centrifuged at 2,000 x g for 20 minutes, followed by 10,000xg for 20 minutes. Then, 0.5 volume of 1X PBS was added followed by 0.05 volume of proteinase K and the sample was incubated at 37°C for 10 minutes. The exosomes precipitation reagent was then added at 0.2 volume, incubate at 4°C for 30 minutes and centrifuged at 10,000 x g for 30 minutes at 4°C. The correct vesicle diameter size for exosomes (∼100nm) was confirmed by Zetaview particle analysis see **Suppl Fig. 1**.

### 2.7 Brain dissection and exosome extraction from brain

Animals were anesthetized with isoflurane, then the brain was then extracted and immediately placed in a solution of ice-cold sterile phosphate-buffered saline (PBS). The prefrontal cortex (PFC), hippocampus, amygdala, and somatosensory cortex were then dissected according to the Paxinos atlas [21]. For exosome extraction, each brain region from four animals within the same experimental group was combined to ensure a representative pool. Exosomes were isolated from brain tissue as described [22]. Tissue was first digested with an enzyme solution containing 2 mg/ml collagenase in Hibernate E for 15 minutes at a temperature of 37°C. After digestion, a protease/phosphatase inhibitor was added in a volume equal to two times that of the total sample volume. The mixture underwent sequential centrifugations at specific speeds (300g for 10 minutes, 2,000g for 10 minutes, and 10,000g for 25 minutes) at 4°C. The desired exosome-containing supernatant was carefully collected. To further refine the exosome isolation, the collected supernatant was passed through a 0.45 μm filter. The exosomes were subsequently extracted using qEVorigincal 35mm Gen2 columns (Izon Sciences). Specific fractions (7-9) from the column elution were collected, concentrating the exosome samples using a 100 kDa cut-off tube. The correct vesicle diameter size for exosomes (∼100nm) was confirmed by Zetaview particle analysis see **Suppl Fig. 1**.

### 2.8 Isolation of Astrocyte Exosome by Immunoprecipitation

Astrocyte-derived exosomes (A-Exo) were purified from total exosome fractions from the blood and brain as described [23]. Briefly, samples were incubated with 3µg anti-GLAST (ACSA-1) biotinylated antibody (Miltenyi Biotec, USA) and BSA (3%) for 4 hours at 4°C after which streptavidin-agarose UltraLink Resin (Thermo Fisher Scientific) for 1 hour at 4°C. After a brief centrifugation (200g, 10 min, 4°C), the supernatant and pellet were used for RNA extraction.

### 2.9 Primary astrocyte culture and exosome purification

Our published procedure was used[24] Briefly, primary astrocyte cultures, were derived from P0-P3 mouse brain. Exosomes were prepared from 72hrs astrocyte conditioned medium (ACM) that was serum free. ACM was first spun at 300 × g for 10 min at room temperature to remove suspension cells, then at 2000 × g for 10 min at 4 °C to remove cell debris, then underwent following purification steps or stored at −80 °C. ACM supernatant was first concentrated (to 500 μl) by centrifugation at 3500 × g for 30 min at 4 °C using Centricon® Plus-70 Centrifugal Filter Devices with a 30k molecular weight cutoff (MilliporeSigma). Then the concentrated supernatant was passed through a 0.22 μm PES filter. The qEVoriginal 35 nm columns Gen2 (Izon Science, MA, USA) were then used as described above. Finally, the purity of exosomes was confirmed by tunable resistive pulse sensing (see **Suppl. Fig 2**).

### 2.10 Astrocyte-derived exosome injection

Astrocyte-derived exosomes (10ug protein) or saline controls were injected in the tail vein at day D0, D3 and D5 in a final volume of 100µl.

### 2.11 RNA extraction, cDNA synthesis and Real-time PCR

Total RNA, including miRNA, was extracted from A-Exos and the remaining exosomes using single-cell RNA extraction kit (Norgen Biotek, Canada), according to manufacturer protocol. Concentration of RNA was determined using a NanoDrop-1000 (Thermofisher Scientific). 10 ng of RNA from each sample was reverse transcribed using TaqMan Advanced miRNA cDNA Synthesis Kit (ABI, USA). Real-time PCR was performed for each target and sample in duplicate on a StepOnePlus PCR System (Applied Biosystems) with primers from TaqMan™ Advanced miRNA Assay. All data were analyzed using the Comparative ΔΔCT method to generate relative expression data using an endogenous miRNA that does not change under conditions used in these experiments as the internal control for miRNAs, as is standard procedure [25]. **Suppl. Fig. 3** confirms that miR-192 is a suitable internal control for these experiments as it was for our previous study on CSI stress and miR-34c [15].

### 2.12 Protein Extraction and Western Blotting

Protein extraction from exosomes was carried out using RIPA lysis buffer containing 50 mM Tris-base at pH 7.5, 0.1% sodium dodecyl sulfate, 0.5% sodium deoxycholate, and 150 mM sodium chloride. The protein concentration was determined using NanoDrop spectrophotometry. Electrophoresis was carried out on a 10% polyacrylamide gel, and the separated proteins were transferred onto a polyvinylidene fluoride membrane (Bio-Rad, USA). Overnight incubation at 4°C with a primary antibody (GLAST1, CD81, TSG101 and CD63ABclonal, USA) at a 1/1000 dilution in skim milk followed by a one-hour incubation with a secondary antibody (Goat anti-Rabbit, ABclonal, USA, 1/3000) at room temperature. The blots were then washed three times with TBST, and visualization of protein bands was achieved using enhanced chemiluminescence (ECL, Bio-Rad, USA) in Chemidoc (Bio-Rad, USA).

### 2.13 Experimental Design and Statistical analysis

Real-time PCR data were analyzed using ExpressionSuite Software (v1.3), which employs the ΔΔCt (delta-delta Ct) method for relative quantification of gene expression. A p-value less than 0.05 was considered significant.

## 3 RESULTS

### 3.1 Cauda sperm and epididymosome and levels of miR-34c are suppressed by CSI stress

Various sperm non-coding RNAs, including miRNAs, are incorporated into sperm via RNA-carrying exosomes secreted by epithelial cells of the cauda (final) section of the epididymis, called epididymosomes. The RNA content of sperm can be altered when the environment changes epididymosome RNA content [26]. To investigate how CSI stress controls sperm miRNA levels we assessed the relevance of this pathway. **Fig. 1** shows that it is, since while miR-34c levels are unchanged by CSI stress in sperm that just exited the testes and entered the caput epididymis **(Fig.1A)**, they are reduced 5-6-fold in sperm (p=0.0365) **(Fig. 1B)** and epididymosomes **(Fig. 1C)** purified from the cauda epididymis (p=0.0489).

**Figure 1.**
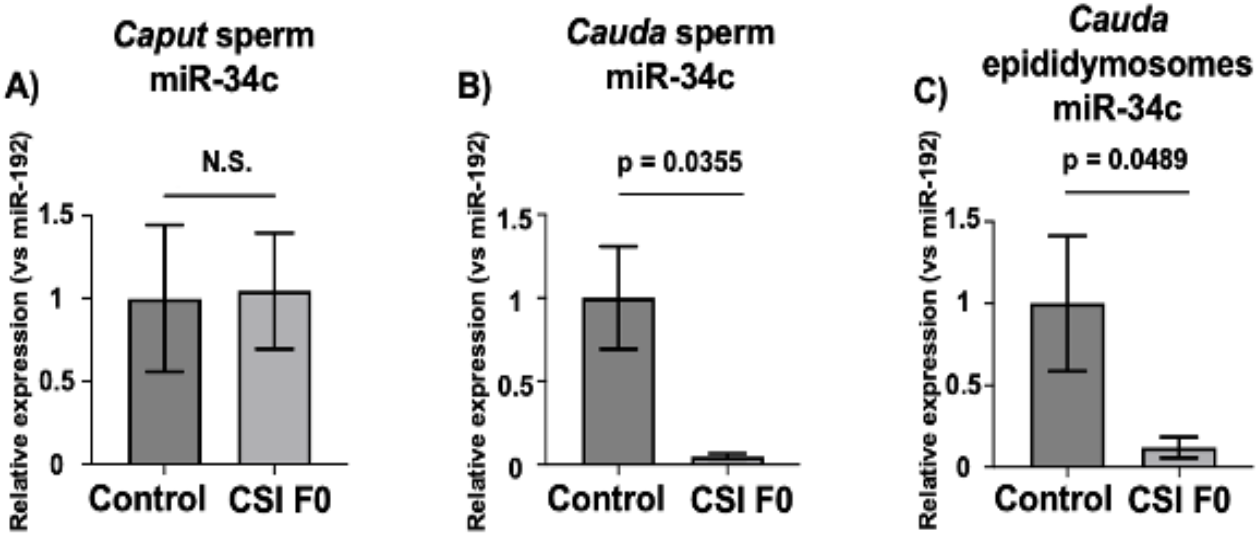
CSI stress reduces miR-34c levels in epididymosomes and sperm isolated from the *cauda* epididymis, but not sperm isolated from the *caput* epididymis. miR-34c levels in sperm isolated from *caput* **(A)** and *cauda* **(B)** regions of the epididymis from control and CSI stressed males. **C)** miR-34c levels in epididymosomes isolated from the *cauda* epididymis of control and CSI stressed males. n=5 for *caput* and *cauda* sperm and n=4 for *cauda* epididymosomes. Data are expressed as mean ± S.E.M; Mann-Whitney test, Two-tailed, WT: wild-type, CSI: Chronic Social Instability. Each point represents an individual mouse. All values are compared to the internal control miR-192 that does not change enough by any perturbation to significantly alter the conclusions (see **Suppl. Fig. 4**) and control ratios are set to 1.

### 3.2 CSI stress suppresses blood astrocyte-derived exosome content of miR-34c across generations

The notion that a specific signal from the brain to germ cells may contribute to the transmission of acquired traits to offspring is supported by studies in *c. elegans* that demonstrated neurons can send RNA signals to germ cells for this purpose[27]. That it could be derived from astrocytes was suggested by the fact that astrocytes are highly responsive to stress and the proof of principle study in mammals showing that when a human miRNA was expressed specifically in brain neurons and astrocytes of mice by viral infection, the human miRNA was subsequently found in the epididymis of males and offspring after their mating [28]. The possibility that the stress sensitive signal could be miR-34c itself is suggested by the fact that miRNAs are known to be transported in blood by exosomes, and miR-34c has already been detected in them [29].

We began by testing **a**strocyte-derived **exos**omes (**A-Exos**.) using a well-characterized protocol to purify them from the total pool of isolated blood exosomes. This was achieved through immunoprecipitation with antibodies to the astrocyte-specific surface marker Glast1 [23]. **Fig. 2A** shows that this method effectively isolated A-Exos, as evidenced by as immunoprecipitation by co-immunoprecipition of the exosome marker CD81, and omitting the Glast1 antibody led to undetectable levels of CD81 on the beads. miR-34c was also undetectable (data not shown). **Fig. 2B** reveals that miR-34c levels in A-Exos in the blood of CSI-stressed males were ∼5-fold lower (p=0.0389) than that found in A-Exos from control mice. Importantly, a similar, if not greater, decline in miR-34c A-Exos content (p=0.0055) was found in the F1 male offspring of CSI stressed males, who also exhibited reduced miR-34c in their sperm. In contrast, miR-375, which increases in sperm after exposure of males to various forms of chronic stress that produce different effects in offspring [10],[11] remained unchanged across generations (**Fig. 2C**). Interestingly, no significant change was observed in blood A-Exos miR-34c content in F1 female offspring of CSI stressed males (see **Suppl Fig. 4**). This observation is consistent with our initial observation that transmission through the female lineage fades across generations, suggesting its transmission does not occur through the germline[12].

**Figure 2.**
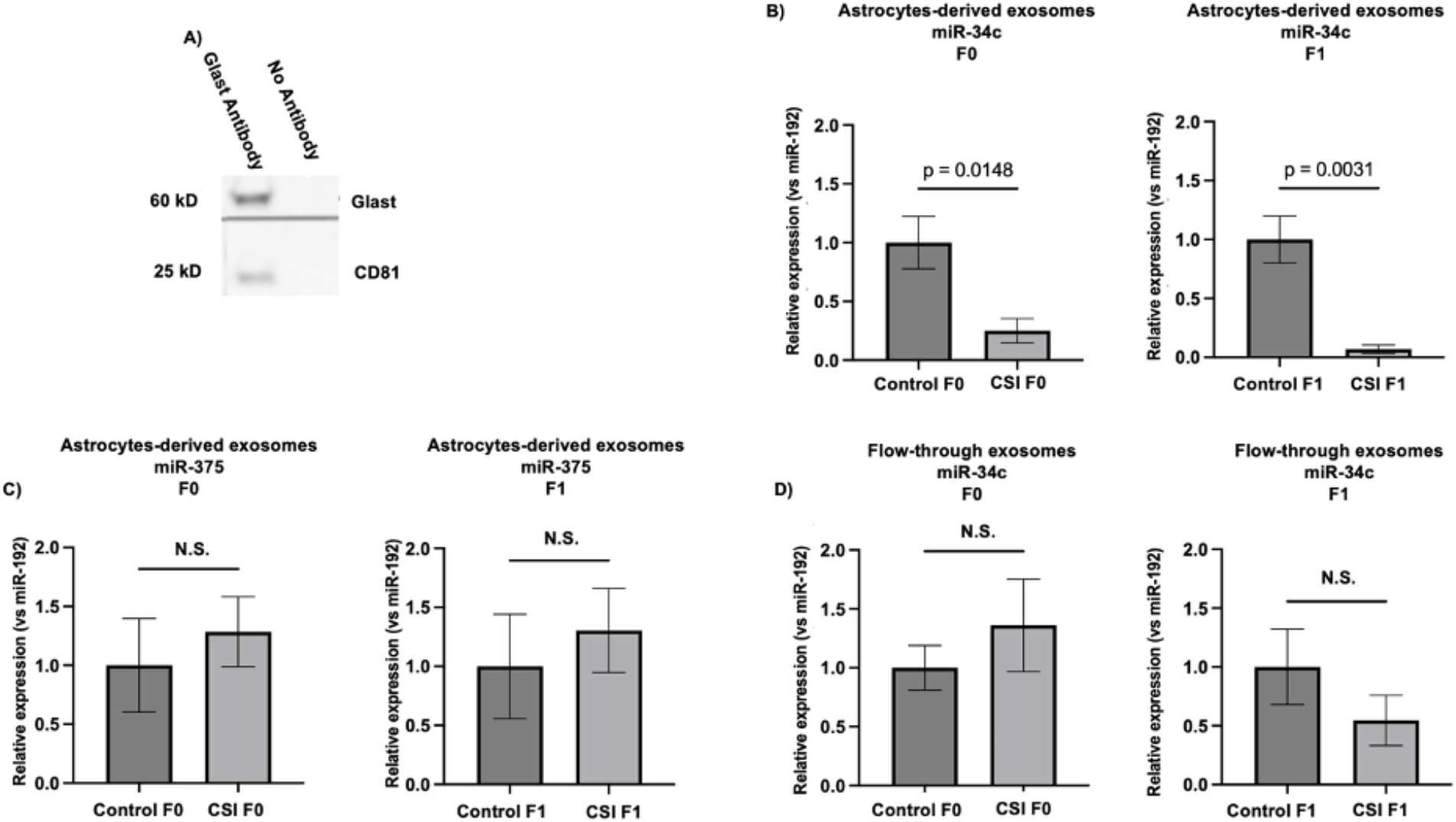
CSI stress suppresses the content of miR-34c in male, blood astrocyte-derived exosomes across generations. **A)** Co-IP of CD81 on western blot after immunoprecipitation with either anti-Glast antibody (Ab) or without (No Ab). **B)** Levels of miR-34c in astrocytes-derived exosomes immunoprecipitated from total plasma exosomes from CSI stress F0 males (left) and their F1 (right) male offspring. N=7 for control and CSI F0 and N=8 for CSI F1 **C)** Levels of miR-375 in astrocytes-derived exosomes isolated from plasma of CSI stressed males (left) and their F1 male offspring (right). N=7 for control and CSI F0 and N=8 for CSI F1; **D)** Levels of miR-34c in the non-precipitated flow-through after immunoprecipitation of astrocytes-derived exosomes. N=6 for control CSI F0 (left) and CSI F1 (right). Data are expressed as mean ± S.E.M; Unpaired t-test with Welch’s correction. WT: wild-type, CSI: Chronic Social Instability. All values are compared to the internal control miR-192 that does not change enough by any perturbation to significantly alter the conclusions (see **Suppl. Fig. 4)** and control ratios are set to 1.

miR-34c was also detectable in the non-precipitated fraction of blood exosomes, which contains exosomes from other cell types in the brain and peripheral tissues. However, its level in this fraction was not affected by CSI stress (**Fig. 2D**). This finding supports the notion that the miR-34c signal between the brain and sperm predominantly exists in A-Exos. CSI stress appears to reduce miR-34c content without altering the number of secreted A-Exos because levels of A-Exos miR-375 (and the internal control miRNA, miR-192-see **Suppl. Fig 4**) do not differ in control and stressed fractions. However, if only a small fraction of A-Exos in the blood carry significant amounts of stress-responsive miR-34c, the reduced levels of that fraction of exosomes could be masked by the exosomes not affected by stres .

In contrast, while miR-449a is also suppressed in sperm of CSI stressed mice and their male F1 offspring, it was not detectable in blood A-Exos (data not shown). This suggests that the regulation of miR-449a in sperm may differ from that of miR-34c. However, our recent findings indicate that reduced sperm miR-34c levels are sufficient to transmit traits to offspring, because when we transiently rescued miR-34c levels in preimplantation embryos derived from CSI stresses males, miR-449a levels were also restored. This occurred by sperm-derived miR-34c increasing its expression from the miR-449 gene in early embryos [15]. Therefore, if a similar mechanism occurs in the epididymis, suppression of miR-34c containing A-Exos might also lead to reduced levels of miR-449 in epididymosomes and subsequently in sperm. This topic is presently under investigation.

### 3.3 Reduced levels of A-Exos miR-34c levels in blood of CSI stressed males are responsible for the reduced levels of miR-34c in sperm

To demonstrate that the CSI-induced suppression of miR-34c levels in blood A-Exos is a critical factor in regulating sperm miR-34c levels, we elevated miR-34c levels in blood exosomes of stressed males by intravenous injection of miR-34c-containing A-Exos purified from cultured mouse astrocytes (as described in [30]), and then tested whether sperm miR-34c levels were restored. The feasibility of testing the biological activity of miR-34c in A-Exos by intravenous injection was supported previous studies, which demonstrated that such injections showing could protect against brain injury induced by cerebral ischemia/reperfusion injury [29].

Mouse sperm remain in the cauda epididymis for up to a week, where they continuously absorb epididymosomes. To determine the optimal number of A-Exos injections required to normalize blood levels in CSI stressed males, we conducted pilot experiments. We decided administering 1ug exosome protein on days 0, 3 and 5 (**Fig. 3A)**. Then, we repeated the experiment either saline or A-Exos injections and measured blood levels of miR-34c on days 4 and 7, as well as epididymosome and sperm miR-34c levels on day 7. Results in **Fig. 3B-D** indicate that restoring miR-34c levels in the blood also led to restoration in cauda epididymosomes and sperm. This finding confirms that the level of miR-34c in blood A-Exos regulates its level in sperm.

**Figure 3.**
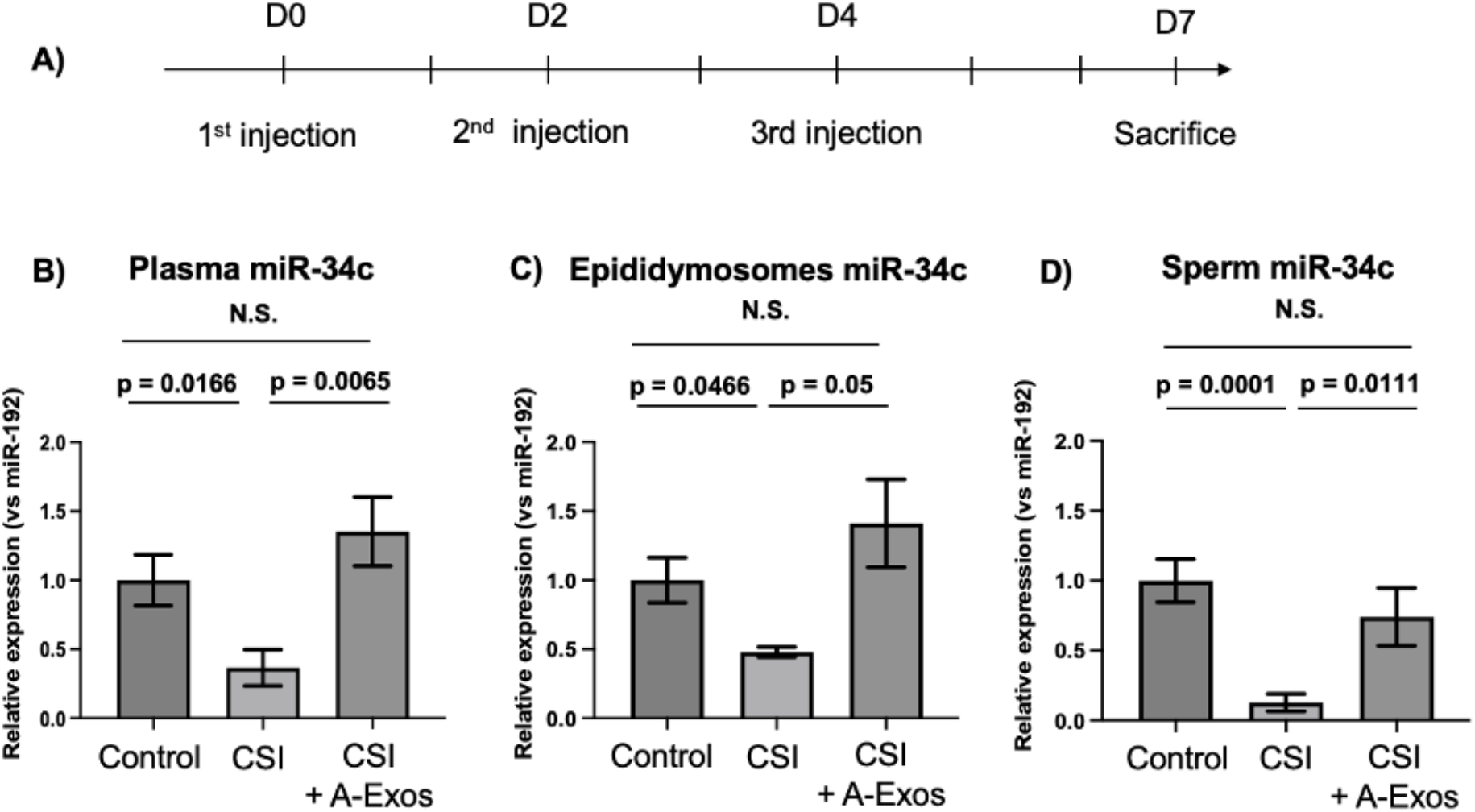
Elevation of blood levels of A-Exos miR-34c in CSI stressed males restores sperm levels of miR-34c. **A)** Scheme for injection of astrocytes-derived exosomes or saline into CSI stressed males and blood collection. **B)** Plasma levels of miR-34c of unstressed (control) males, and the average plasma levels of miR-34c sampled at day 4 and 7 from saline injected (CSI) or exosome injected (CSI+A-Exos) CSI stressed mice; N=7 for unstressed and N=4 for CSI stressed mice. **C)** Levels of miR-34c in epididymosomes isolated from unstressed males (control), and CSI stressed males injected with either saline (CSI) or purified astrocyte-derived exosomes (CSI + A-Exos) Welch’s T-test was used. N= 4 for control, saline and exosome injected mice. **D)** The same as C except miR-34c levels were assayed in cauda sperm; Welch’s T-test was used. N= 12 for control and N=4 for saline or exosome injected mice CSI stressed mice. Data are expressed as mean ± S.E.M; CSI: Chronic Social Instability. Each point represents an individual mouse. All values are compared to the internal control miR-192 that does not change enough by any perturbation to significantly alter the conclusions (see **Suppl. Fig. 4** and control ratios are set to 1.

### 3.4 CSI stress reduces level of A-Exos miR-34c in the pre-frontal cortex and amygdala, but not hippocampus, of both stressed males and their F1 male offspring

To reveal brain regions contributing to the depletion of miR-34c-bearing A-Exos in the blood of CSI-stressed males and their F1 offspring, we measured miR-34c levels in A-Exos purified from brain regions known to respond to chronic stress, as well as one that is not. We first isolated the total exosome fraction from the adult brain as previously described [22], and then used Glast1 antibodies to specifically immunoprecipitaate the A-Exos fraction. As observed in the blood A-Exos studies (**Fig. 2)**, immunoprecipitation with Glast1 antibodies was specific, as miR-34c levels were undetectable when immunoprecipitations were performed without Glast1 antibodies.

We began with the **p**re**f**rontal **c**ortex **(PFC)**, known to have altered transcription profile in response to CSI stress [31]. **Fig. 4A** shows a ∼20-fold decline in miR-34c levels in A-Exos isolated from both directly stressed mice (p=0.02) and their F1 male offspring (p=0.009). Similar to blood A-Exos, no significant change in miR-375 levels was detected in either A-Exos population for miR-375, despite its elevation in sperm due to different stress paradigms [10], [11]. miR-34c was also detectable in the non-precipitated flow through fraction which contains exosomes from other brain cell types. However, its level in this fraction was not changed by CSI stress exposure in either directly stressed mice nor their F1 male offspring (**Fig. 4B**). This finding supports the model that miR-34c in A-Exos, rather than exosomes from other brain cell types, are the major regulators of CSI stress-induced changes in sperm miR-34c content. Also, miR-449a was not detectable in A-Exos isolated from the PFC (data not shown) consistent with data above on blood A-Exos.

**Figure 4.**
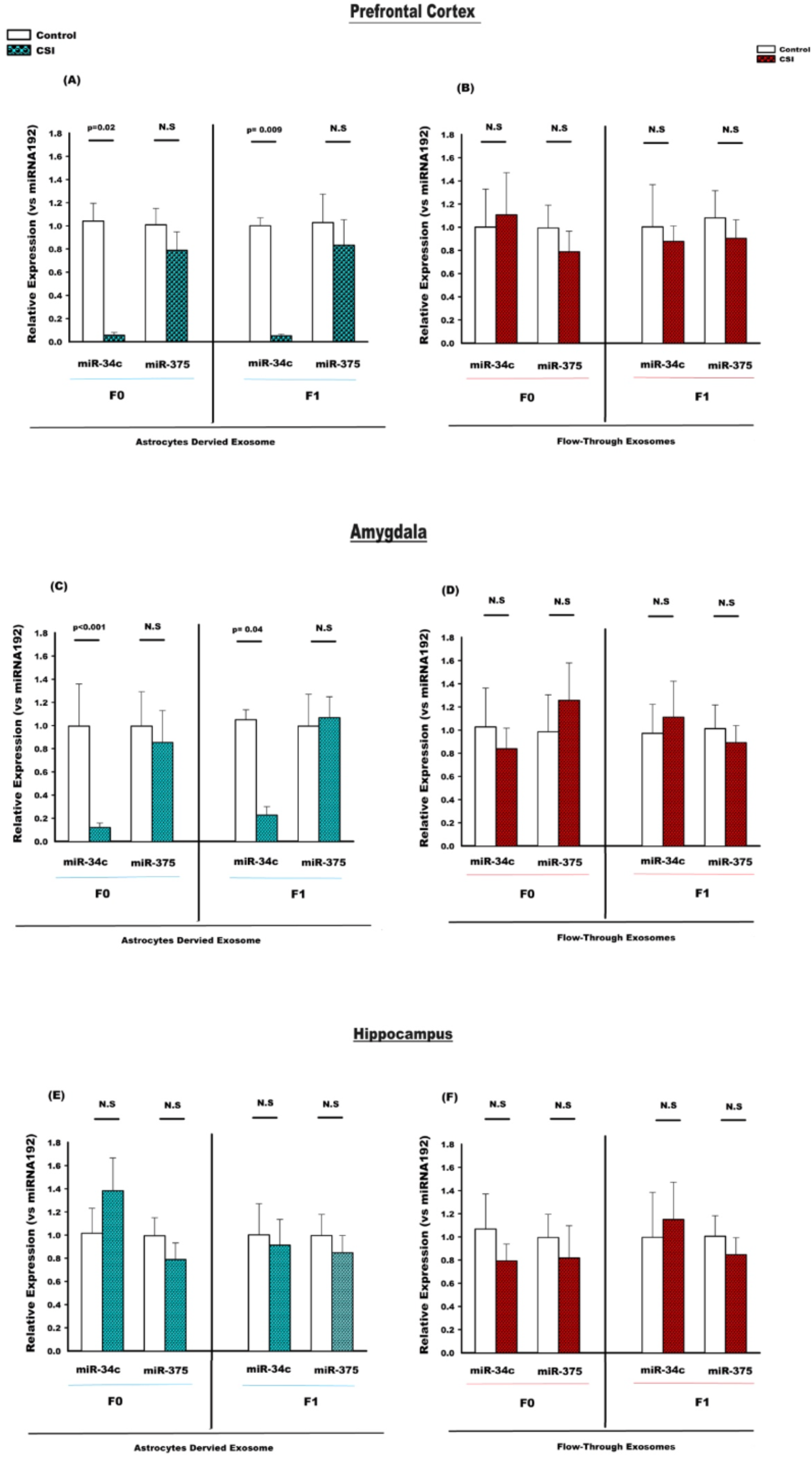
CSI suppresses levels of miR-34c in A-Exos isolated from the prefrontal cortex and amygdala, but not hippocampus, across generations. **A-B) Prefrontal Cortex-A)-Left-** miR-34c and miR-375 levels in A-Exos immune-purified from the total pool of exosomes in the prefrontal cortex of control and CSI-stressed F0 males (n=7), **Right-**the same analysis on A-Exos from their F1 male offspring (n=9 for miR-34c, n=5 for miR-375). **B) Left**-miR-34c and miR-375 levels in the non-precipitated, flow-through fraction of exosomes purified from the prefrontal cortex of control and CSI-stressed F0 males (n=5), Right-the same analysis on exosomes from their F1 male offspring (n=5) **C-D**) **Amygdala-C)-Left-** miR-34c and miR-375 levels in A-Exos immune-purified from the total pool of exosomes in the amygdala of control and CSI stressed males (n=6) **Right-**the same analysis on A-Exos from their F1 male offspring (n=7); **D) Left**-miR-34c and miR-375 levels in the non-precipitated, flow-through fraction of exosomes purified from the amygdala and CSI-stressed F0 males (n=5), Right-the same analysis on exosomes from their F1 male offspring (n=5). **E-F) Hippocampus**-**E) Left-**miR-34c and miR-375 levels in A-Exos immune-purified from the total pool of exosomes in hippocampus of control and CSI-stressed F0 males (n=5), **Right-**the same analysis on A-Exos from their F1 male offspring (n=5). **F) Left**-miR-34c and miR-375 levels in the non-precipitated, flow-through fraction of exosomes purified from the hippocampus of control and CSI-stressed F0 males (n=5), **Right**-the same analysis on exosomes from their F1 male offspring (n=5) . Bars represent the fold differences in mean normalized expression values, and the error bars are standard errors (+/-SEM). Each data point is a pool of tissues from 4 mice. All values are compared to the internal control miR-192 that does not change enough by any perturbation to significantly alter the conclusions and control ratios are set to 1 (see **Suppl. Fig. 4**).

In the amygdala, miR-34c levels in A-Exos were also suppressed by CSI stress, though to a lesser degree (∼5-fold), across generations of males (**Fig. 4C** p<0.001 for F0, and p=0.04 for F1). Similar to findings in the PFC, the non-astrocyte-derived exosome fraction containe significant miR-34c, but its level was also not affected by exposure of mice to CSI stress (**Fig. 4D**).

In contrast to the PFC and amygdala, miR-34c levels were not suppressed in A-Exos isolated from the hippocampus of CSI stressed males, nor in their male offspring (**Fig. 4E**). Similar to the PFC and amygdala, the non-astrocyte-derived exosome fraction in the hippocampus also contained significant levels of miR-34c, but these levels were unaffected by CSI-stress (**Fig. 4F**). This implies that that other types of exosomes in the hippocampus do not contribute to the CSI stress response. Finally, we found that A-Exos isolated from the sensory cortex, a region not known to be stress-responsive, did not contain detectable levels of miR-34c (data not shown).

## 4 DISCUSSION

There is increasing evidence that a significant portin of inherited susceptibility to mental health disorders may stem from one’s parent’s experiences transmitted to them via epigenetic changes in parent’s germ cells [32]. Much of this evidence derives from mouse models of chronic stress, where stress-specific changes in sperm miRNA content leads to corresponding alterations in offspring phenotypes. However, the mechanisms by which specific stressors can alter sperm miRNA content have remained unclear. The experiments described here reveal that CSI stress reduces sperm levels of miR-34c by suppressing the miR-34c content in astrocyte-derived exosomes (**A-Exos)** present in blood, since we demonstrated that restoring blood A-Exos miR-34c content in these mice restored miR-34c levels in their sperm. We implicated the prefrontal cortex and amygdala as sources of miR-34c-containing A-Exos that are suppressed by CSI stress, as the level of A-Exos miR-34c in these brain regions was severely reduced not only in CSI-stressed males but also in their F1 male offspring, both of which display reduced levels of miR-34c in their blood A-Exos and sperm. However, until we can define the specific astrocyte populations in these processes and test the effect of manipulating their exosome function directly, we must rely upon this strong correlative evidence. Interestingly, the hippocampus, another stress-responsive brain region, also contains significant levels of A-Exos miR-34c, but these levels are not affected by CSI stress. This suggests that hippocampal exosomes may serve a more localized role for miR-34c, or mediate transgenerational effects of different stressors not yet reported to function them.

These experiments also revealed the surprising finding that A-Exos are normally responsible for sustaining the level of miR-34c in sperm. CSI stress inhibits miR-34c levels carried by them in the blood, likely by suppressing their release from astrocytes in the prefrontal cortex and amygdala, and possibly other brain regions. As a result, sperm miR-34c levels are suppressed. Additionally, these A-Exos in blood may regulate miR-34c levels in other stress-responsive tissues, and thus be part of a novel mechanism to alter peripheral tissue function in a stress-specific manner.

Several key questions remain to be revealed, including identifying the specific astrocyte populations involved. Also, how does CSI stress alter astrocyte function in the PFC and amygdala, but not hippocampus, such that either less miR-34c is uploaded into each exosome produced or less miR-34c-containing exosomes are generated. CSI stress may suppress cellular miR-34c uploading into existing exosomes by either suppressing cellular miR-34c levels or by inhibiting regulators that upload specific miRNAs into exosomes (for review see [33]). Alternatively, the seven weeks of CSI stress beginning during adolescence may lead to the loss of the specific astrocyte populations in the PFC and amygdala ito explain the large ∼20-fold decline in miR-34c levels. Why this may occur in the PFC but not hippocampus could be due to the difference in their susceptibility to chronic stress during adolescence. In particular, numerous studies [34-37] have demonstrated that stress induces dendritic retractions and synaptic losses in both adolescent and adult medial PFC. Notably, dendritic branching can recover to pre-stress levels when adult subjects are removed from stressful conditions [38]. However, such recoveries are not observed following adolescent social instability stress, suggesting that adverse experiences during adolescence, when the CSI stress we impose begins, may permanently disrupt typical PFC astrocyte development [31]. Interestingly this effect was not observed in the hippocampus [39]. Moreover, there is a significant enhancement in amygdala-PFC connectivity during adolescence [40]. Thus, the changes in A-Exo miR-34c cargo that we observed only in the PFC and amygdala might be attributed to the fact that this connectivity undergoes late maturation, making it more vulnerable compared to other regions like the hippocampus.

Another striking finding is that astrocytes in the same brain regions in F1 offspring of stressed mice “inherit” the same exosome dysfunction observed in their fathers, despite never being exposed to stress themselves. This inherited defect explains the observed reductions in A-Exos miR-34c levels in both the blood and sperm of these offspring, leading to transmission of stress-related traits to subsequent offspring. The inherited phenotype in these astrocytes likely stems from compromised astrocyte development that arose as a consequence of reduced miR-34c during their preimplantation embryo stage, a defect we showed contributes to reduced miR-34/449 levels in their sperm [41].

We also identified miR-409-3p as a sperm miRNA involved in this example of epigenetic inheritance. Its levels increase in sperm from mice exposed to CSI stress and in their male offspring [14]. However, we could not detect miR-409-3p in A-Exos from the PFC or blood (data not shown), suggesting that the regulation of sperm miRNAs whose levels rise in response to chronic stress are regulated by different mechanisms. This idea is supported by our finding that miR-375, which also increases in sperm in response to both maternal separation at birth combined with unpredictable maternal stress and chronic variable stress[10, 11], is present in A-Exos from the PFC and amygdala. But unlike miR-34c, miR-375 level does not change in response to CSI stress. Interestingly, we hypothesize that miRNAs whose sperm levels rise in response to chronic stress function differently at another step in the epigenetic inheritance process; how stress-induced change in their very low content in sperm can have a large impact on early-embryos upon fertilization [41].

These findings open several intriguing areas for future research. For example, there is growing interest in the potential use of brain-derived blood exosome content as a biomarker for neurological disorders [42]. The results presented here suggest that reduced A-Exos miR-34c content in blood A-Exos could indicate early life trauma. This hypothesis is supported by our previous research indicating that miR-34c levels are also reduced in sperm of men who were exposed to a high level of Adverse Childhood Experiences (ACEs) [13], a finding subsequently corroborated using an alternative scale of early life trauma [43].

## Supplementary Figure Legends

**Supplemental Figure 1:**
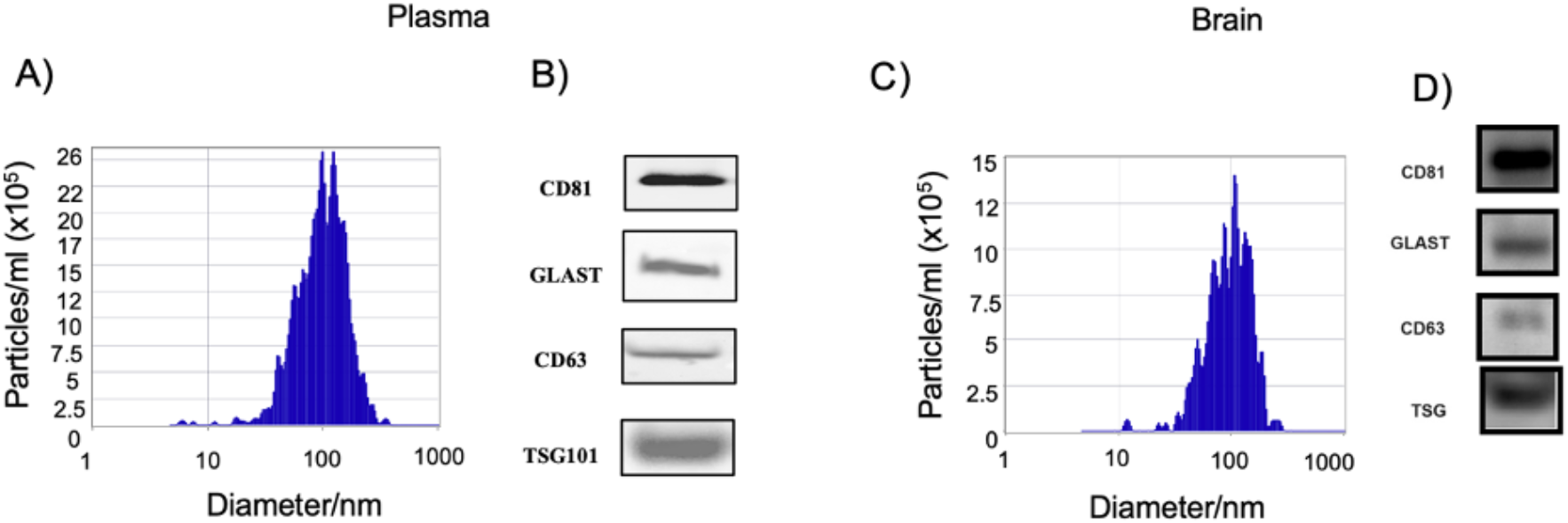
Representative size distribution histogram of exosomes. Total exosomes isolated from plasma and brain from control mouse have been analysed by the Zetaview particle analyser (A, C) and by Western-blot of CD81, Glast, CD63 and TSG markers (B, D).

**Supplemental Figure 2:**
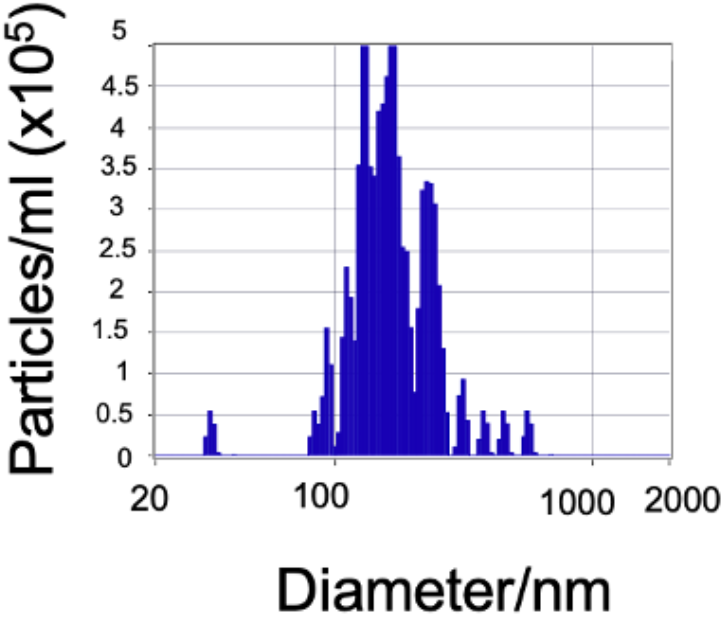
Total exosomes isolated from astrocytes cell culture have been analysed by the Zetaview.

**Supplemental Figure 3:**
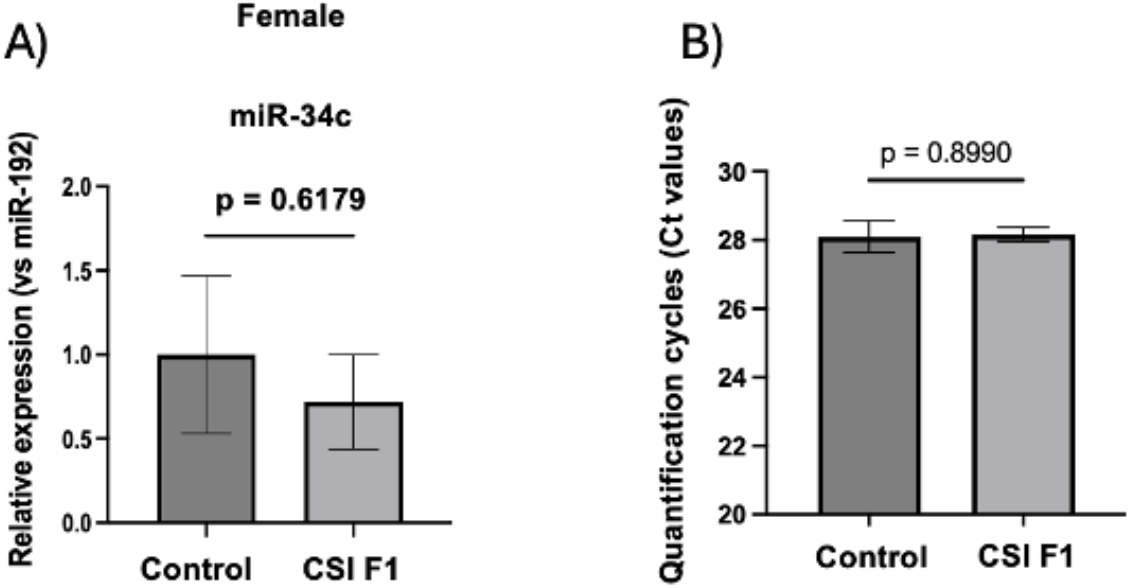
Astrocytes-exosomes miR-34c in blood of F1 CSI females. A) Astrocytes-exosomes (A-Exos) were isolated from plasma of F1 females derived from mating control or CSI stressed males. The level of miR-34c is unchanged in A-Exos. B) Level of miR-192 (Ct value) in A-Exos in blood from F1 females derived from mating control or CSI stressed males. Data are expressed as mean ± S.E.M; T-test. All values are compared to the internal control miR-192 that does not change enough by any perturbation to significantly alter the conclusions and control ratios are set to 1. N=7 for control and N=8 for CSI F1

**Supplemental figure 4:**
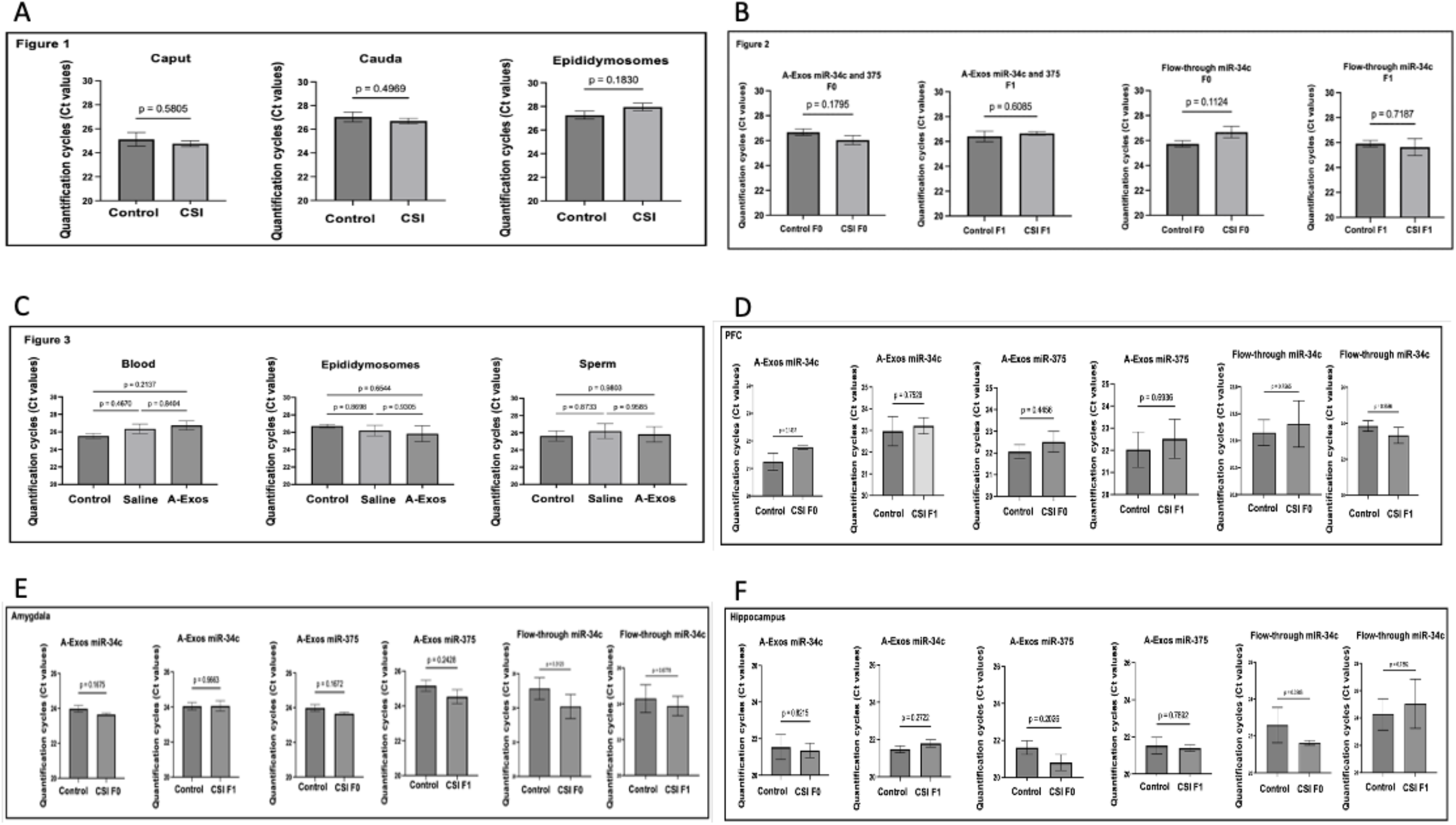
Quantification of cycles of internal normalizer, miR192. A) Quantification cycles of miR-192 from figure 1. B) Quantification cycles of miR-192 from figure 2. C) Quantification cycles of miR-192 from figure 3. D) Quantification cycles of miR-192 from figure 4 on PFC . E) Quantification cycles of miR-192 from figure 4 on Amygdala. F) Quantification cycles of miR-192 from figure 4 on Hippocampus . Data are shown as mean of Ct values +/-SEM. One-way Anova with multiple comparisons for 3 groups (figures 2 and 3) and T-test for figures 1 (A) and 4 (D-F) . PFC: prefrontal cortex.

